# Motile and non-motile *Listeria* species adopt distinct genomic and ecological strategies to achieve broad geographic ranges in soil

**DOI:** 10.64898/2026.01.08.698536

**Authors:** Ying-Xian Goh, Shannon Hepp, Kevin J. Cummings, Martin Wiedmann, Jingqiu Liao

**Author notes:** Corresponding author: Jingqiu Liao, Mailing address: 403 Durham Hall, 1145 Perry Street, Blacksburg, VA 24061.

## Abstract

Broad geographic ranges indicate high ecological versatility and are generally associated with low extinction risk. Motility is not only a physiological feature, but also a core ecological trait for bacteria. However, the genomic foundations underlying the broad geographic ranges of motile and non-motile bacteria remain poorly understood. To address this, we analyzed 141 and 90 genomes of *Listeria welshimeri* and *L. booriae*, systematically screened from 1,004 soil samples across the United States, representing widespread motile and non-motile species, respectively. We show that *L. welshimeri* exhibits restricted phylogeographic structure and a weak distance-decay relationship, suggesting minimal geographic barriers to dispersal. Its high dispersal capacity likely stems from enhanced motility and effective host colonization, evidenced by strong positive selection on flagellar genes and the close relatedness to isolates from wild birds. In contrast, *L. booriae* displays a regionally endemic distribution and limited dispersal. With a large, open pangenome, *L. booriae* appears strongly adapted to local environments. This is evidenced by pronounced positive selection on genes involved in inorganic ion, amino acid, and coenzyme transport and metabolism, as well as strong associations with abiotic factors, particularly climate, and accompanying bacterial consortia. In summary, to establish widespread distributions, *L. welshimeri* tends to rely on movement and colonization of wildlife hosts to facilitate long-distance dispersal, while *L. booriae* leverages genomic plasticity and metabolic versatility that enable adaptation to diverse environmental conditions. These findings highlight the distinct genomic foundations and ecological strategies that motile and non-motile bacteria use to achieve broad geographic ranges in the environment.

## Introduction

Geographic range, the spatial extent over which a species occurs, is a key determinant of microbial ecology, associating with population connectivity, resilience, and ecosystem functioning [1, 2]. Species with broader ranges typically exploit a wider variety of habitats or resources, reflecting greater niche breadth, and are often considered generalists [3–5]. Although geographic range expansion can entail fitness costs, such as reduced performance in the original habitat due to niche-expanding mutations [6] or slower adaptation to novel conditions [7], generalists generally gain a competitive advantage under environmental fluctuations, making them less prone to extinction when habitats are disturbed [8, 9]. Yet microbial communities are often dominated by specialists [10–12], raising the question of why some taxa achieve broad geographic distribution while others remain narrowly restricted. While evolutionary processes such as pangenome expansion [13] and recombination [14] have been proposed to facilitate bacterial survival across diverse environments, large-scale field data that mechanistically link genomic features to their widespread geographic distributions remain limited.

Free-living bacteria can be broadly categorized into motile and non-motile forms. Motility confers multiple ecological advantages, including microhabitat exploration, efficient nutrient acquisition, toxin avoidance, habitat and host colonization, and dispersal across locations [15–17]. Experimental studies show that motile bacteria expand at rates determined by growth and habitat size [18], with chemotaxis enabling directed range expansion [19]. However, these benefits are counterbalanced by substantial energetic costs, including the biosynthesis and maintenance of flagellar structures and ATP expenditure for locomotion [15]. Trade-offs between chemotactic benefits and energetic burden may constrain the net advantage of motility, particularly in complex or turbulent environments [20], leaving open the question of whether mechanistic advantages observed under laboratory conditions translate into broad-scale geographic distribution in natural habitats. In contrast, non-motile bacteria lack active dispersal and may rely on passive mechanisms, such as rain [21], wind [22, 23], or intermicrobial hitchhiking [24, 25], which are likely less effective in penetrating structured environments such as soil due to physical fragmentation of pathways. This suggests that non-motile taxa may evolve alternative strategies to achieve broad geographic ranges, which remain poorly understood.

*Listeria* is a bacterial genus widespread across diverse environments [26, 27]. Despite sharing a common evolutionary background, we previously found that *Listeria* species have undergone adaptive evolution, showing pronounced differences in geographic distribution in soil [28]. Notably, we observed that *L. welshimeri*, a species that is motile via peritrichous flagella [29], and *L. booriae*, a non-motile species [30], are both widely distributed in soil across the contiguous United States (US) [28]. The contrasting motility phenotypes but comparable geographic range sizes make these two *Listeria* species an ideal model for investigating the mechanisms underlying ability of motile and non-motile bacteria to disperse to and occupy the broad geographic ranges.

Here, we performed an in-depth comparative genomic and ecological analysis of 141 *L. welshimeri* and 90 *L. booriae* isolates obtained from our systematic screening of 1,004 soil samples across the US [28]. Using phylogeographic, bioinformatic, and statistical approaches, we found that flagellar-associated genes in *L. welshimeri* show evidence of positive selection, and that this species is capable of long-distance dispersal, as indicated by a weak distance-decay relationship. To test whether wildlife may facilitate this dispersal, we performed whole-genome sequencing (WGS) of nine *L. welshimeri* isolates recovered from wild birds and compared them with soil isolates. Their high genomic relatedness supports a potential role for wildlife as a transport vector that enable long-distance dispersal. In contrast, *L. booriae*, which exhibited strong dispersal constraints, appears to offset the absence of motility by maintaining an extremely open pangenome enriched in adaptive signatures related to metabolism, allowing them to exploit a wide range of environmental niches. Indeed, using a suite of statistical analyses, we identified strong correlations between the gene content of *L. booriae* and abiotic factors, particularly climate, as well as accompanying bacterial consortia, particularly Proteobacteria, Chloroflexi, and Actinobacteria, suggesting genetic adaption to local environments. Together, these findings demonstrate that motile and non-motile bacteria achieve widespread geographic distribution in soil through distinct genomic and ecological strategies, offering broader insights into microbial adaptation and persistence in heterogeneous environments.

## Materials and Methods

### Genomic data of *L. welshimeri* and *L. booriae* isolates, environmental data, and bacterial community data

Existing genome assemblies of 141 *L. welshimeri* and 90 *L. booriae* isolates that we previously obtained from minimally disturbed soil in natural environments across the U.S. [28] were subjected to secondary analysis in this study. The genome assemblies of these two species were previously analyzed along with other species in the context of *Listeria* pangenome evolution but have not been used to directly compare the genomic features between each other. Soil sampling followed a systematic approach that achieved a relatively even spatial coverage across the U.S., and the methods for DNA extraction, WGS and data preprocessing, genome assembly, gene prediction, ortholog gene identification, and functional annotation (i.e., assignment of functional categories for ortholog genes based on Clusters of Orthologous Groups [COGs] and Kyoto Encyclopedia of Genes and Genomes [KEGG]) for these soil isolates were detailed in Liao et al. (2021) [28].

Additionally, WGS was performed for nine *L. welshimeri* isolates recovered from wild birds, including Rose-breasted Grosbeak and Rock Pigeon, admitted to the Janet L. Swanson Wildlife Hospital at Cornell University, New York [31]. Following our data deposition, these isolates remained the only *L. welshimeri* wildlife isolates with WGS data available in the NCBI database. Genomic DNA was extracted using the NEBExpress T4 Lysozyme and Monarch Spin gDNA Extraction Kit (New England Biolabs) following the manufacturer protocol. DNA quality and concentration were assessed using a NanoDrop spectrophotometer and Qubit fluorometer, respectively. All samples met standard quality thresholds (A260/280 ≈ 1.80; A260/230 ≈ 2.0) and were sequenced on the Illumina MiSeq platform (2 × 250 bp paired-end reads) at the Duke University Center for Genomic and Computational Biology. WGS data preprocessing, genome assembly, gene prediction and gene family identification followed the workflow described in Liao et al. (2021) [28] with minor modification to ensure comparability. In brief, sequencing adapters and low-quality reads were trimmed using fastp v0.23.4 [32], and cleaned reads were assembled de novo with SPAdes v4.0.0 [33]. Contigs <500 bp were removed. Assembly quality was assessed with QUAST v5.2.0 [34] and CheckM2 v1.0.2 [35], and average sequencing depth was estimated using BBmap v39.01 and Samtools v1.17. All assemblies met the following criteria: <300 contigs, N50 >50,000 bp, average coverage >30×, completeness >90%, contamination <5%, and correct taxonomic assignment by Kraken2 v2.1.3 [36]. Quality statistics for wild bird isolates are summarized in **Supplementary Table 1**. Genes of wild bird isolates were predicted using Prodigal v2.6.3 [37], and ortholog genes present in both soil and wild bird isolates were identified using MMseqs2 [38].

The environmental dataset (also referred to as “abiotic factors”) for soil isolates comprised 34 variables: three geolocation variables (latitude, longitude, elevation), 17 soil physicochemical variables (moisture, total nitrogen [TN], total carbon [TC], pH, organic matter [OM], and concentrations of aluminum [Al], calcium [Ca], copper [Cu], iron [Fe], potassium [K], magnesium [Mg], manganese [Mn], molybdenum [Mo], sodium [Na], phosphorous [P], sulfur [S], and zinc [Zn]), four climate (precipitation, wind speed, maximum [Max.] and minimum [Max.] temperatures [temp]), and 10 land use variables (proportional coverage of open water, barren land, forest, shrubland, grassland, cropland, pasture, wetland, and developed [Dev] open spaces with either >20% or <20% impervious [IMP] surfaces in the surrounding landscape). Descriptions of environmental data acquisition and processing were detailed in Liao et al. (2021) [28].

Bacterial community data (also referred to as “biotic factors”) for soil isolates, including methods for DNA extraction, 16S rRNA gene amplicon sequencing, sequencing data preprocessing, operational taxonomic unit (OTU) identification, taxonomic classification, and diversity calculation (e.g., weighted UniFrac distances) were reported in our previous study comparing *Listeria* populations from natural and food-associated environments [39]. Both abiotic and biotic factors were subjected to secondary analysis in this present study to assess their differential associations with *L. welshimeri* and *L. booriae* as well as their associations with gene content.

### Phylogenetic trees, genetic similarities, and geographic distribution

To investigate phylogeographic structure, phylogenetic trees based on core SNPs were constructed for three groups: (i) *L. welshimeri* soil isolates, (ii) *L. booriae* soil isolates, and (iii) *L. welshimeri* soil and wild bird isolates. Core SNPs were identified using kSNP4 [40], and maximum likelihood trees were generated using RAxML v8.2.12 [41] under the GTR + G + I substitution model with ascertainment bias correction and 1,000 bootstrap replicates. Trees were visualized using iTOL v7 [42]. To assess genomic similarity, pairwise average nucleotide identity (ANI) was computed using pyani v0.3.0-alpha in Python v3.6.8 for these three groups. In addition, Jaccard similarity indices were calculated using the gene presence/absence matrices for groups (i) and (ii).

Geographic distributions of *L. welshimeri* and *L. booriae* soil isolates grouped by major clades on the phylogenetic tree were visualized using a Mercator projection with the Basemap Matplotlib Toolkit v1.2.1 in Python v3.6.8. Pairwise geographic distances between isolates were calculated from GPS coordinates using the geopy module in Python v3.6.8. Mean geographic distances within each major clade were then computed and compared between the two species using a two-sided Mann-Whitney (MW) *U* test.

### Pangenome characterization and statistical comparison of genomic features

In this study, core genes are defined as those found in all genomes of a given species, whereas accessory genes are defined as those present in only a subset of genomes [13, 43, 44]. The pangenome refers to the complete repertoire of genes present across all genomes within each species, encompassing both core and accessory genes [13, 45]. To estimate the pangenome size for soil-derived *L. welshimeri* and *L. booriae* based on an assumed 100 genomes each, we applied the power law function *cN^γ^* calculated in Liao et al. (2021) [28], which models the relationship between the number of genomes (*N*) and total ortholog genes (i.e., pangenome size). The specific functions used were n_pan_ = 2572*N*^0.133^ for *L. welshimeri* and n_pan_ = 2937*N*^0.208^ for *L. booriae* [28].

To characterize genomic differences between soil-derived *L. welshimeri* and *L. booriae*, we analyzed genome size, gene richness (i.e., total number of unique ortholog genes per genome), nucleotide diversity (π), relative abundance of COGs, relative abundance of putatively functional motility and chemotaxis genes, and KEGG pathway diversity. Species-level π was calculated using popGenStat [46] on gene alignments generated with MUSCLE v3.8.31 [47]. Reference sequences for 31 motility and five chemotaxis genes (**Supplementary Table 2**) were retrieved from the BIGSdb-*Lm* database [48] and used as BLASTN queries against assembled genomes to identify homologs [49]. For each gene, the highest bit-score hit was screened for premature stop codons (excluding terminal codon: TGA, TAG, TAA) and alignment coverage in Python 3.6.8. Motility and chemotaxis genes with >80% coverage and no premature stops were considered putatively functional [39]. Genes were further classified into KEGG pathways and analyzed for classification into four major pathways (i.e., metabolism, genetic information processing, environmental information processing, cellular processes; **Supplementary Table 3**). Shannon-Wiener diversity index was calculated for all pathways combined (referred to as “overall biological pathway”) and within each major pathway based on their relative abundance. All genomic features were compared between *L. welshimeri* and *L. booriae* soil isolates using two-sided MW *U* tests. For the comparison of COG relative abundance, Benjamini-Hochberg false discovery rate (BH-FDR) correction was additionally applied to account for multiple testing.

To assess the genomic similarities between monophyletic *L. welshimeri* soil and wild bird isolates, Kruskal-Wallis (KW) tests followed by pairwise two-sided MW *U* tests were used to compare ANI for (i) within-soil, (ii) soil-wild bird, and (iii) within-wild bird isolates. Genome size, gene richness, and GC content were also compared using two-sided MW *U* tests.

### Positive selection detection and functional enrichment analysis

Soil isolates of *L. welshimeri* and *L. booriae* were subjected to positive selection detection using the methods detailed in Liao et al. (2021) [28]. Briefly, core and accessory genes of each species were screened for positive selection using the branch-site unrestricted statistical test for episodic diversification (BUSTED) model implemented in Hypothesis Testing Using Phylogenies (HyPhy) [50]. Genes were included if they contained at least three non-identical sequences and exhibited one or more non-synonymous substitutions. Evidence of positive selection was evaluated using a likelihood ratio test (LRT) comparing unconstrained and null models, with significance assessed via a χ^2^ test followed by BH-FDR correction. Homologous recombination was also assessed for both core and accessory genes using the Recombination Detection Program v4 (RDP4) [51]. Only genes with an adjusted *P* < 0.05 in the positive selection analysis and no detectable recombination were considered undergoing positive selection. Functional enrichment analysis was performed using a binomial distribution model, as described in Liao et al. 2023 [39], to identify COGs significantly enriched among core and accessory genes under positive selection for each species.

### Environmental characteristics and distance-decay relationships

Multidimensional scaling (MDS) analyses was performed using Euclidean distances for abiotic factors and weighted UniFrac distances for OTUs to compare environmental abiotic conditions and bacterial community compositions between soil samples positive for *L. welshimeri* and *L. booriae*. Clustering significance by species was assessed with PERMANOVA. Differences in individual abiotic and biotic variables between species were evaluated using two-sided MW *U* tests with BH-FDR correction.

To identify the influence of abiotic and biotic factors on the gene richness of soil-derived *L. welshimeri* and *L. booriae*, variation partitioning analysis (VPA) and Spearman’s rank correlation analyses were conducted. Two VPA models were constructed: (i) one that included three groups of abiotic variables - soil properties, climate, and surrounding land use - and (ii) another that included two groups of variables, abiotic factors and biotic factors represented by relative abundances of bacterial phyla. To enhance model performance, we evaluated multicollinearity among predictor variables using variance inflation factors (VIF). Variables with the highest VIF were iteratively removed until all remaining predictors had VIF < 10 [52]. This procedure retained 22 and 42 variables for VPA models (i) and (ii), respectively (**Supplementary Table 4**), to quantify the proportion of variance in gene richness explained by each group of selected variables within each species. Adjusted *R^2^* values were computed using the vegan package 2.6-4 in R and visualized in a Venn diagram. Spearman’s rank correlations followed by BH-FDR correction were applied to assess associations between gene richness, both overall and for each COG, and each of the 34 abiotic factors or 25 bacterial phyla.

To infer dispersal limitation, distance-decay relationships were examined by regressing geographic distance and two measures of genomic similarity: ANI and Jaccard similarity. A linear regression was interpreted as evidence of a distance-decay relationship when it met all three criteria: an *R^2^* > 0.2, a negative slope, and *P* < 0.05.

## Results

### Soil-derived *L. welshimeri* and *L. booriae* exhibits contrasting phylogeography and genomic features

Our previous work showed that *L. welshimeri* and *L. booriae* were the most common motile and non-motile *Listeria* species detected in systematically collected, minimally disturbed natural soils across the contiguous U.S. [28]. Among 1,004 soil samples, 89 (8.86%) and 67 (6.67%) were positive for *L. welshimeri* and *L. booriae*, respectively [28]. While *L. monocytogenes* is also motile and exhibited a slightly higher overall prevalence (11.75%) than *L. welshimeri*, our previous systematic phylogenomic analysis revealed that *L. monocytogenes* comprises three species-level phylogroups (lineages I, II, and III) which substantially differ in genomic and ecological traits, whereas *L. welshimeri* consists of a single phylogroup [28, 53]. At the same phylogenetic clustering level, *L. welshimeri* was more frequent than the most common *L. monocytogenes* lineage (III; 7.67%) and was therefore selected to represent the most widespread motile *Listeria* species in this study. Both *L. welshimeri* and *L. booriae* exhibited comparable broad geographic distributions (853 km and 850 km, respectively; **Fig. 1A-B**). Core-SNP phylogenies revealed that *L. welshimeri* can be classified into nine major clades (**Fig. 1C**), whose members were geographically intermixed (**Fig. 1A**). *L. booriae* also comprised nine major clades (**Fig. 1D**) but exhibited clear geographic structuring, with major clades endemic to certain regions (**Fig. 1B**). To support this observation, pairwise geographic distances between isolates within each major clade were calculated for each species. The mean geographic distances within major clades ranged from 234-963 km and 33-730 km for *L. welshimeri* and *L. booriae*, respectively (**Supplementary Fig. 1A**) and were significantly larger in *L. welshimeri* (MW *U P* = 0.022; **Supplementary Fig. 1B**). These findings indicate that although both *L. welshimeri* and *L. booriae* are widespread, they show distinct geographic and population structures.

**Figure 1.**
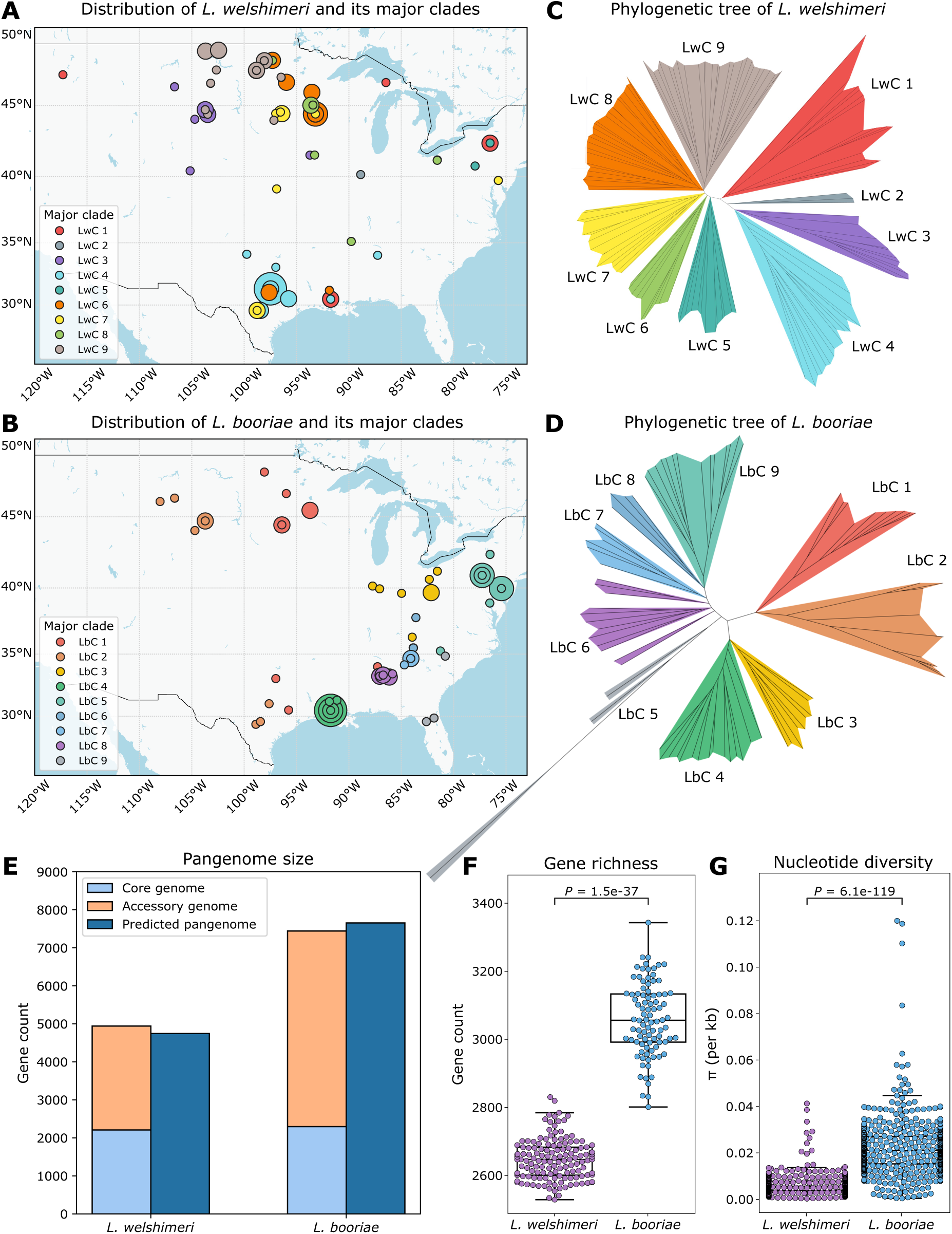
Phylogeographic structure and genomic features of soil-dwelling *L. welshimeri* and *L. booriae.* **(A-B)** Geographic distribution of **(A)** *L. welshimeri* and **(B)** *L. booriae* isolates in soil across the US, color-coded by major clades identified in **C-D**. Circle sizes indicate the number of isolates per clade at each site. **(C-D)** Unrooted maximum likelihood phylogenetic trees of **(C)** 141 *L. welshimeri* soil isolates and **(D)** 90 *L. booriae* soil isolates based on core SNPs. Branches are color-coded by major clades, with “Lw” and “Lb” indicating *L. welshimeri* and *L. booriae*, respectively. **(E)** Pangenome sizes of *L. welshimeri* and *L. booriae*, divided into core and accessory genomes, with predicted pangenome sizes estimated from 100 genomes per species using the power law function *cN^γ^* from Liao et al. (2021) [28]. **(F-G) (F)** Gene richness and **(G)** nucleotide diversity (π) compared between *L. welshimeri* and *L. booriae*. Box plots display the interquartile range (IQR) with the median indicated as a line and whiskers extending to 1.5 times the IQR. Two-sided MW *U P* values are annotated.

To assess whether the observed population structures corresponded to differences in genomic content between the two species, we performed pangenome analysis, which identified 4,941 ortholog genes in *L. welshimeri* (2,209 core and 2,732 accessory genes) and 7,443 in *L. booriae* (2,298 core and 5,145 accessory genes) (**Fig. 1E**). To avoid the potential bias caused by unequal sample size, we predicted the pangenome size given 100 genomes for each species based on the pangenome curves, where a more open pangenome was observed in *L. booriae* (**Supplementary Fig. 2A**). The predicted pangenome size was 4,745 for *L. welshimeri* and 7,654 for *L. booriae.* Consistent with this result, *L. booriae* had significantly higher gene richness (MW *U P* = 1.5e-37; **Fig. 1F**) and larger genome size (*P* = 1.5e-37; **Supplementary Fig. 2B**). Also, it had significantly greater nucleotide diversity (*P* = 6.1e-199; **Fig. 1G**). Together, these results show that *L. booriae* has greater genomic diversity and plasticity than *L. welshimeri*.

### Soil-derived *L. welshimeri* and *L. booriae* undergo strong positive selection targeting flagellar genes and metabolic genes, respectively

To assess the genomic differences between *L. welshimeri* and *L. booriae* reflected at the functional level, we analyzed their COG functional profiles. Eleven COGs, including cell motility (N) and replication, recombination, and repair (L), were significantly more abundant in *L. welshimeri*, whereas nine, including cell membrane biogenesis (M), were enriched in *L. booriae* (adjusted MW *U P* < 0.05 for all; **Fig. 2A**). Among these, cell motility (N) genes showed the largest fold change in *L. welshimeri* (adjusted *P* = 5.4e-37; **Fig 2A**), and targeted BLAST searches confirmed significantly more putatively functional motility and chemotaxis genes in this species (*P* = 9.8e-40 and 1.5e-37, respectively; **Supplementary Fig. 3, Fig. 2B**). In contrast, cell membrane biogenesis (M) genes exhibited the largest fold change in *L. booriae*, being significantly more abundant in this species (adjusted *P* = 7.4e-37; **Fig 2A**). These results reveal distinct functional repertoires between *L. welshimeri* and *L. booriae*.

**Figure 2.**
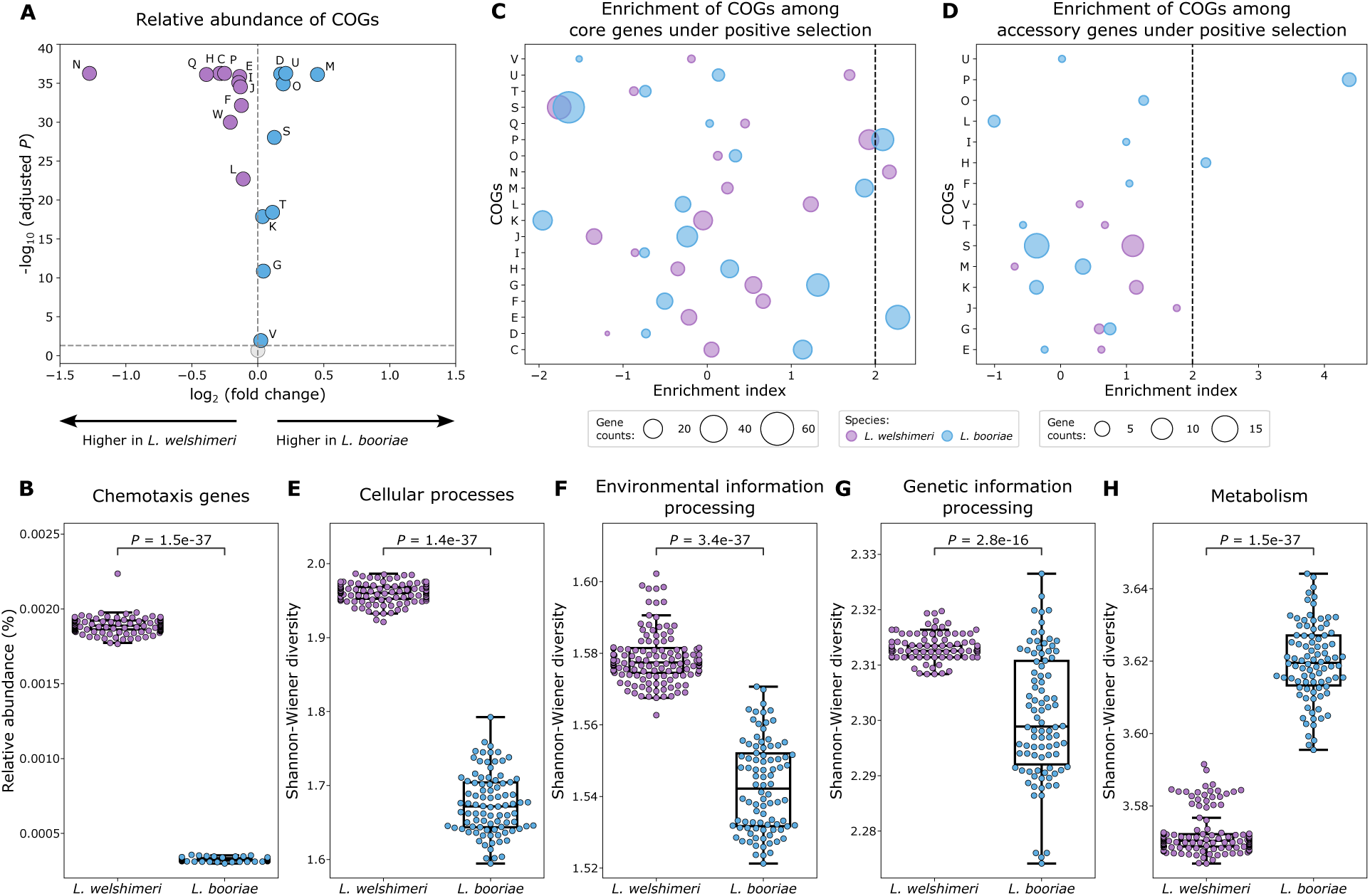
Functional divergence and positive selection in *L. welshimeri* and *L. booriae*. **(A)** Volcano plot of COG abundance differences between *L. welshimeri* and *L. booriae*, with fold change on the x-axis and significance (two-sided MW *U P*) on the y-axis. Points above the horizontal grey dashed line have adjusted *P* < 0.05. **(B)** Relative abundance of putatively functional chemotaxis genes compared between *L. welshimeri* and *L. booriae.* **(C-D)** Enrichment of COGs among **(C)** core and **(D)** accessory genes with evidence of positive selection. An enrichment index > 2 (black dashed line) indicates significant enrichment (binomial *P* < 0.05). Circle size is proportional to the number of genes annotated to each COG. For **(A), (C),** and **(D),** COGs abbreviations are as follows: C: Energy production and conversion; D: Cell cycle control, cell division, chromosome partitioning; E: Amino acid transport and metabolism; F: Nucleotide transport and metabolism; G: Carbohydrate transport and metabolism; H: Coenzyme transport and metabolism; I: Lipid transport and metabolism; J: Translation, ribosomal structure, and biogenesis; K: Transcription; L: Replication, recombination, and repair; M: Cell wall/membrane/envelope biogenesis; N: Cell motility; O: Posttranslational modification, protein turnover, chaperones; P: Inorganic ion transport and metabolism; Q: Secondary metabolites biosynthesis, transport, and catabolism; T: Signal transduction mechanisms; U: Intracellular trafficking, secretion, and vesicular transport; V: Defense mechanisms.**(E-H)** Shannon-Wiener diversity of KEGG pathways compared between *L. welshimeri* and *L. booriae* for **(E)** cellular processes, **(F)** environmental information processing, **(G)** genetic information processing, and **(H)** metabolism pathways. For **(B), (E-H),** box plots display the IQR with the median indicated as a line and whiskers extending to 1.5 times the IQR. Two-sided MW *U P* values are annotated.

Because niche breadth in *Listeria* has been linked to positive selection [28], we next identified genes under positive selection in each species using the BUSTED model for gene-wide episodic selection. In *L. welshimeri*, 193 core and 31 accessory genes exhibited evidence of positive selection (adjusted LRT *P* < 0.05 for all; **Supplementary Table 5**). Based on the functional enrichment analysis, core genes of *L. welshimeri* under positive selection were significantly enriched for cell motility (N) functions (*P* < 0.05; **Fig. 2C**), including a chemotaxis protein and several flagellar synthesis genes (e.g., *fliH, fliK, flhB, flgC, fliM*; **Supplementary Table 5**). No COGs were significantly enriched among accessory genes under positive selection in this species (**Fig. 2D**). By contrast, *L. booriae* displayed a broader set of positively selected genes, which included 290 core and 64 accessory genes (adjusted LRT *P* < 0.05 for all; **Supplementary Table 5**). Core genes under positive selection in this species were enriched for inorganic ion transport and metabolism (P) and amino acid transport and metabolism (E) (**Fig. 2C**), while accessory genes under positive selection were enriched for inorganic ion transport and metabolism (P) and coenzyme transport and metabolism (H) (**Figs. 2D**). Approximately half of the positively selected ion transport genes encoded ATP-binding cassette (ABC) transporters, localized to the cell membrane (**Supplementary Table 5**). These patterns indicate that positive selection has primarily shaped motility capacity in *L. welshimeri* and metabolic versatility in *L. booriae*.

To determine whether the functional differences extended to higher-level gene networks, we analyzed KEGG pathway diversity. *L. welshimeri* exhibited significantly higher pathway diversity in cellular processes (MW *U P* = 1.4e-37; **Fig. 2E**) and environmental information processing (*P* = 3.4e-37; **Fig. 2F**). This increased diversity may reflect the presence of flagellar assembly pathways and an expanded repertoire of sensing and regulatory systems (e.g., phosphotransferase system and two-component systems). *L. welshimeri* also showed significantly higher pathway diversity in genetic information processing (*P* = 2.8e-16; **Fig. 2G**), which includes essential modules involved in DNA replication, recombination, and repair (**Supplementary Table 3**). In contrast, *L. booriae* displayed significantly greater diversity in both metabolism and overall biological pathways (*P* = 1.5e-37 and 4.5e-14, respectively; **Fig. 2H, Supplementary Fig. 4**), indicating broader metabolic potential to thrive in diverse local environments. These findings reveal distinct genomic functional specialization between *L. welshimeri* and *L. booriae*.

### *L. booriae* shows strong genomic associations with soil environments

Because major clades of *L. booriae* exhibited endemism (**Fig. 1B**) and this species possessed a flexible, adaptive pangenome (**Fig. 1E, Supplementary Fig. 2A**), we hypothesized that its genome content is strongly shaped by local environmental conditions, facilitating adaptation to diverse soil habitats and enabling its broad geographic distribution. To test this hypothesis, we compared the abiotic conditions and bacterial communities of *L. welshimeri* and *L. booriae* habitats and assessed their correlations with gene richness of each species. A total of 21 abiotic factors and 15 bacterial phyla were found to be significantly different between their habitats (adjusted MW *U P* < 0.05 for all). *L. welshimeri* was primarily isolated from sites with significantly higher minimum temperature, elevation, pH, and concentration of certain soil macronutrients (K, Mg, Ca, P), greater coverage of pasture, wetland, and cropland in surrounding areas (**Supplementary Fig. 5A**), and higher relative abundances of Proteobacteria, Planctomycetes, Elusimicrobia, and Chlamydiae (**Supplementary Fig. 5B)**. In contrast, *L. booriae* occurred in sites with significantly higher precipitation, greater forest and shrubland coverage in surrounding areas, elevated soil concentrations of Fe, Al, Zn, Mn, and Cu (**Supplementary Fig. 5A**), and enriched with Actinobacteria, Chloroflexi, Bacteroidetes, Firmicutes, and Fibrobacteres (**Supplementary Fig. 5B**) Consistent with these results, both MDS plots based on Euclidian distance of 34 abiotic variables and bacterial community beta diversity indicated by weighted UniFrac distances revealed significant clustering of samples by species (PERMANOVA *P* = 0.001 for both, pseudo-*F* = 18.84 and 32.60, respectively; **Supplementary Fig. 6A and B**), indicating that *L. welshimeri* and *L. booriae* occupied distinct environmental abiotic and biotic spaces.

For the genomic associations with abiotic factors, VPA using VIF-selected variables (see Methods) showed that overall abiotic factors explained more gene richness variation in *L. booriae* (16.76%) than *L. welshimeri* (12.52%; **Fig. 3A**). Climate and soil properties contributed the largest unique fraction of explained variation in gene richness in *L. booriae* (9.29%) and *L. welshimeri* (10.99%), respectively (**Fig. 3A**). When stratified by COGs, abiotic factors explained more variation in *L. booriae* than in *L. welshimeri* for 15 of 19 COGs, with the strongest effects observed for cell membrane biogenesis (M; 77.93%), followed by signal transduction mechanism (T; 66.54%) and transcription (K; 47.54%; **Fig 3B**). Consistent with the VPA results, Spearman correlation analyses showed that gene richnesses of *L. booriae* exhibited stronger associations with abiotic factors. Overall gene richness in *L. booriae* was positively associated with precipitation and temperature, and negatively with latitude and elevation (adjusted *P* < 0.05 for all; **Fig. 3C**), whereas no significant association was observed for *L. welshimeri* (**Supplementary Fig. 7A**). Across COGs, an average of ten and five abiotic factors were significantly associated with gene richness in *L. booriae* and *L. welshimeri*, respectively (**Fig. 3C; Supplementary Fig. 7A**). Notably, precipitation showed strong correlations (|Spearman’s *ρ*| > 0.5) with four COGs, including inorganic ion transport and metabolism (P) and amino acid transport and metabolism (E; **Fig. 3C**), both of which also exhibited evidence of positive selection in *L. booriae* (**Fig. 2C and D**). Cell membrane biogenesis (M) and posttranslational modification, protein turnover, chaperones (O) displayed the highest number of significant associations with abiotic factors (22 significant variables; **Fig. 3C**), indicating that genes involved in these functions may be particularly responsive to abiotic variation in *L. booriae*. Collectively, these findings indicate that abiotic factors, particularly climate, exert a stronger influence on gene content in *L. booriae* than *L. welshimeri*.

**Figure 3.**
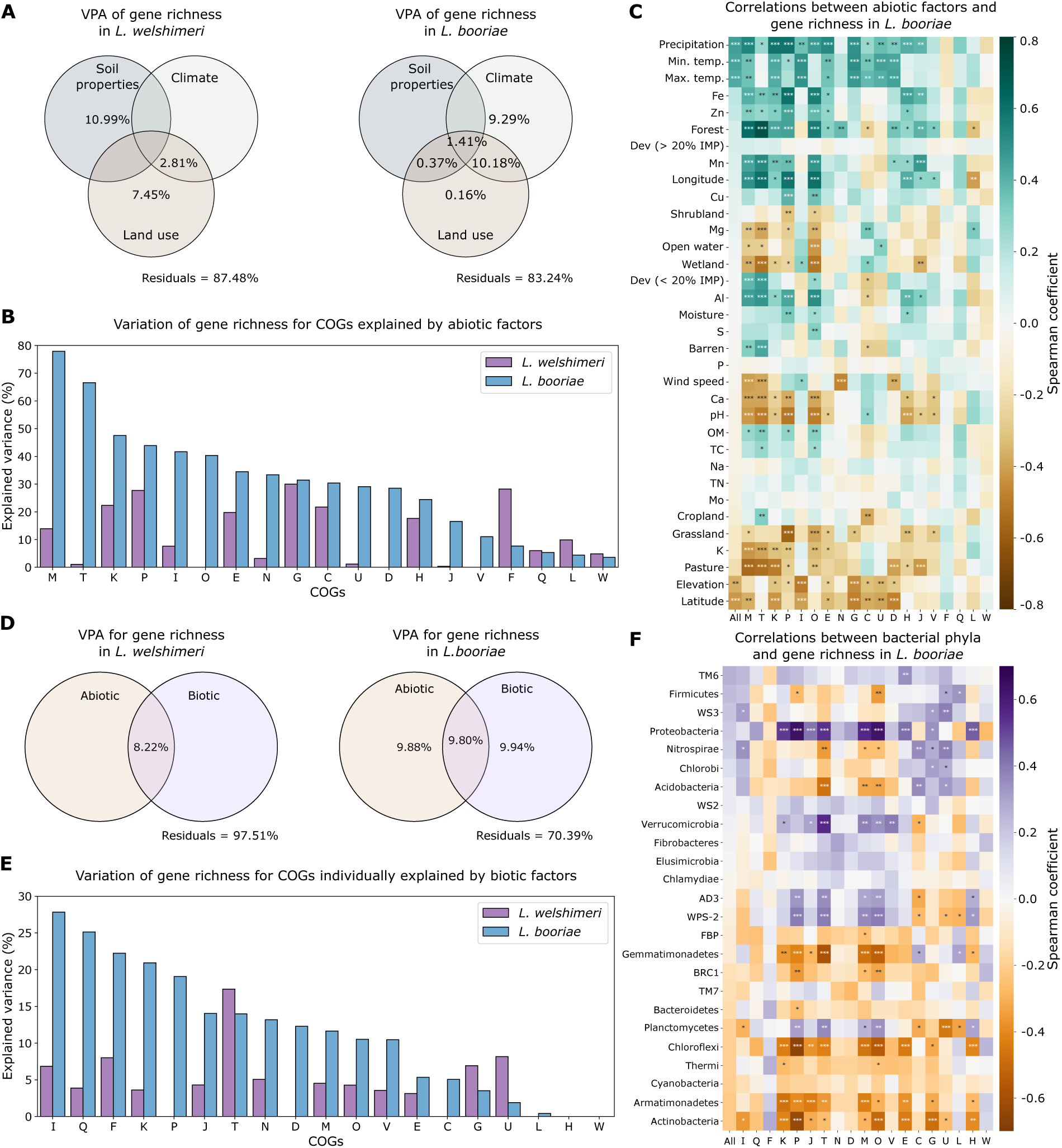
Associations between gene richness in *L. welshimeri* and *L. booriae* and abiotic environmental factors and bacterial community composition. **(A)** Venn diagrams from variation partitioning analysis (VPA) showing the proportion of overall gene richness variation explained by soil properties, climate, and surrounding land use for *L. welshimeri* and *L. booriae*. **(B)** Variation in gene richness of COG categories explained by abiotic factors based on VPA. **(C)** Spearman’s correlations between abiotic factors and gene richness, both overall and for COGs, in *L. booriae*. Abiotic factors are ordered by the descending correlation coefficients of overall (“All”) gene richness. Positive and negative correlations are shown in green and brown, respectively. **(D)** Venn diagrams from VPA showing the proportion of gene richness variation explained by abiotic and biotic factors (i.e., surrounding bacterial phyla) for *L. welshimeri* and *L. booriae.* For **(A)** and **(D)**, only positive fractions of explained variance are shown; negative fractions were omitted, and therefore the total explained variance and the residual does not sum to 1. **(E)** Variation in gene richness of COG categories individually explained by biotic factors based on VPA. **(F)** Spearman’s correlations between the relative abundance of bacterial phyla and gene richness, both overall and for COGs, in *L. booriae*. Bacterial phyla are ordered by the descending correlation coefficients of overall (“All”) gene richness. TM6, WS3, WS2, AD3, WPS-2, FBP, BRC1, and TM7 represent candidate bacterial phyla that remain uncultured under laboratory conditions. Positive and negative correlations are shown in purple and orange, respectively. For **(C)** and **(F)**, significance levels are indicated as “*”, “**”, and “***” for adjusted *P* < 0.05, < 0.01, and < 0.001, respectively.

Additionally, VPA using VIF-selected variables (see Methods) showed that abiotic factors and surrounding bacterial phyla together explained substantially more gene richness variation in *L. booriae* (29.61%) than *L. welshimeri* (2.49%) (**Fig. 3D**). Surrounding bacterial phyla uniquely accounted for 9.94% of the variation in *L. booriae*, while they had minimal explanatory power in *L. welshimeri* (**Fig. 3D**). Notably, in *L. welshimeri*, the total explained variance for overall gene richness decreased from 12.52% (abiotic factors only) to 2.49% when bacterial phyla were included, indicating that bacterial phyla explained little additional variance and that their inclusion resulted in a penalty for increased model complexity [54]. When stratified by COGs, surrounding bacterial phyla individually explained more variation in *L. booriae* than in *L. welshimeri* for 14 of 19 COGs, with the strongest effects observed for lipid transport and metabolism (I; 27.82%), followed by secondary metabolites biosynthesis, transport and catabolism (Q; 25.14%) and nucleotide transport and metabolism (F; 22.25%; **Fig 3E**). Consistent with the VPA results, Spearman correlation analyses indicated that gene richnesses of *L. booriae* exhibited stronger associations with the relative abundance of bacterial phyla. Although overall gene richness in *L. booriae* showed no significant correlations with any individual bacterial phylum (**Fig. 3F**), stratification by COG revealed substantially more significant associations in this species, with an average of six correlations per phylum, compared with an average of one in *L. welshimeri* (**Fig. 3F; Supplementary Fig. 7B**). In *L. booriae*, Actinobacteria, Proteobacteria, and Chloroflexi showed the highest number of strong correlations with COG-specific gene richness (|Spearman’s *ρ*| > 0.5; **Fig. 3F**). Notably, genes involved in inorganic ion transport and metabolism (P), which exhibited signatures of positive selection (**Fig. 2C and D**), were consistently associated with all three phyla in *L. booriae* (**Fig. 3F**). Consistent with patterns observed for abiotic environmental factors (**Fig. 3C**), genes involved in cell membrane biogenesis (M) and posttranslational modification, protein turnover, chaperones (O) exhibited the highest number of significant associations with phyla (13 and 14, respectively; **Fig. 3F**), indicating their key roles in mediating species-community interactions in *L. booriae*. Collectively, these findings indicate that like abiotic factors, bacterial community composition, particularly Proteobacteria, Chloroflexi, and Actinobacteria, is also more important to the gene content of *L. booriae* than *L. welshimeri*.

### Wildlife appears to facilitate long-distance dispersal of *L. welshimeri* in soil

As *L. welshimeri* had minimal geographically confined clades and was not strongly restricted by environmental conditions, we hypothesized that it frequently disperses across geographic locations, facilitating its broad geographic range. To test this hypothesis, we analyzed the distance-decay relationships for both species. The slope of the linear relationship between ANI and geographic distance were 5.5-fold shallower in *L. welshimeri* (*R^2^* = 0.15, slope = -8.79e-07) compared with *L. booriae* (*R^2^* = 0.30, slope = -4.86e-06; **Fig. 4A**), and gene content similarity showed a similar pattern (3.4-fold shallower slopes; *L. welshimeri*: *R^2^* = 0.05, slope = -7.98e-06; *L. booriae*: *R^2^* = 0.22, slope = -2.73e-05; **Supplementary Fig. 8**). These results showed that *L. welshimeri* exhibited a much weaker distance-decay relationship than *L. booriae* and support our hypothesis of frequent dispersal across locations in *L. welshimeri*.

**Figure 4.**
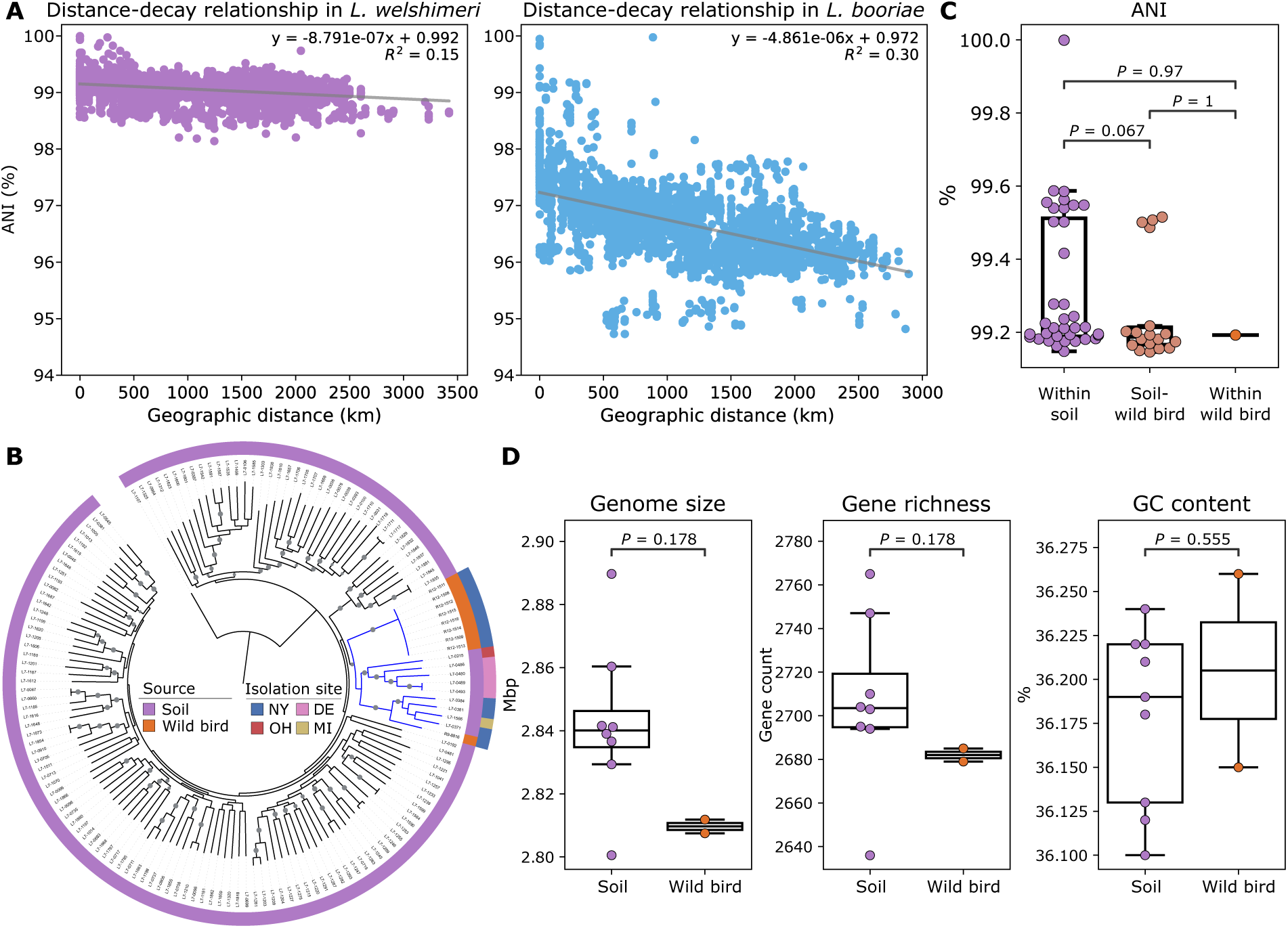
Dispersal patterns of *L. welshimeri* and *L. booriae* and evidence of wild birds as a dispersal vehicle for *L. welshimeri* in soil. **(A)** Distance-decay relationship in *L. welshimeri* and *L. booriae* inferred by the linear regression for genetic similarities measured by average nucleotide identity (ANI) and geographic distances. A steeper negative slope with a higher *R^2^* indicates a stronger distance-decay relationship. **(B)** Maximum likelihood phylogenetic tree based on core SNPs of 141 soil *L. welshimeri* isolates (purple) and nine wild bird isolates (orange). The tree was constructed using 1,000 bootstrap replicates, and bootstrap values > 80% are indicated by gray circles on the corresponding branches. The tree was rooted by midpoint. Branches belonging to the monophyletic clade that contains both soil and wild bird isolates are highlighted in blue, and the collection sites for isolates within this clade are annotated. **(C)** Pairwise ANI comparisons between soil and wild bird isolates of *L. welshimeri*. **(D)** Genome size, gene richness, and GC content compared between monophyletic soil and wild bird *L. welshimeri* isolates. For **(C)** and **(D)**, box plots display the interquartile range (IQR) with the median indicated as a line and whiskers extending to 1.5 times the IQR. Adjusted two-sided MW *U P* values are annotated.

As the reported motility speeds for *Listeria* (∼0.09 μm/s in cytoplasm [55, 56] and ∼10 μm/s on non-charged microscope slide [57]) combined with the physical constraints of soil environments [58, 59] make long-distance dispersal by motility alone improbable, we hypothesized that *L. welshimeri* achieves this via mobile hosts, such as wildlife. To test this hypothesis, we performed WGS of nine *L. welshimeri* isolates recovered from wild birds in New York and compared them with soil isolates. Core-SNP phylogenetic analysis revealed that the wild bird isolates clustered within a monophyletic clade alongside nine soil isolates from nearby regions, including Ohio, Delaware, New York, and Michigan (**Fig. 4B**). Eight of the nine wild bird isolates formed a tight clade with minimal SNPs, suggesting they represent a single strain. R12-1511 was selected as the representative isolate for this strain alongside the other strain R9-8816 and were included in downstream analyses. The representative wild bird and soil isolates forming a monophyletic clade shared high ANI (mean: 99.25%, range: 99.15%-99.52%), with no significant difference observed among within-soil, soil-wild bird, and within-wild bird groups (KW *P* = 0.065; **Fig. 4C**). Comparisons of genome size, gene richness, and GC content also revealed no significant differences between soil and wild bird isolates within the monophyletic clade (MW *U P* = 0.178, 0.178, and 0.555, respectively; **Fig. 4D**). These results indicate that *L. welshimeri* isolates from soil and wild birds are closely related, supporting the hypothesis that wildlife acts as vehicles for long-distance dispersal of *L. welshimeri*.

## Discussion

Here, we show that motile and non-motile *Listeria* species adopt distinct genomic and ecological strategies to achieve broad geographic ranges in soil (**Fig. 5**). *L. welshimeri,* which is a motile species, appears to achieve a widespread distribution through effective long-distance dispersal mediated by wildlife vectors, enabled by enhanced motility and host colonization. In contrast, *L. booriae*, a non-motile species, appears to have limited dispersal and likely attains a broad geographic range through effective environmental adaptation by maintaining an extremely flexible pangenome responsive to abiotic (e.g., climate) and surrounding bacterial phyla (e.g., Proteobacteria, Chloroflexi, and Actinobacteria). This genomic plasticity support fitness primarily through membrane modifications and metabolic versatility, including inorganic ion and amino acid transport and metabolism.

**Figure 5.**
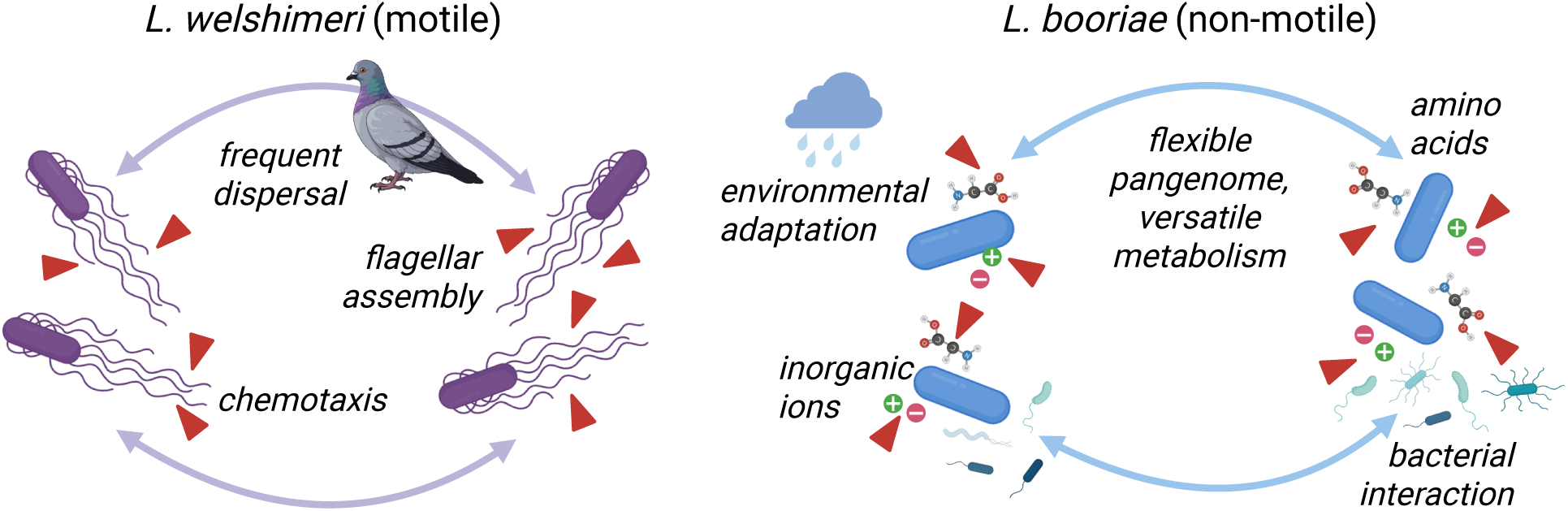
Genomic strategies underlying broad geographic ranges in motile and non-motile *Listeria* species. Triangles mark COG functions targeted by positive selection. Created with BioRender.com.

Flagellar motility and chemotaxis are central to bacterial nutrient sensing and the exploitation of heterogeneous environments [19, 60–63]. In soil, where nutrients are highly patchy and often transient [59], the ability to detect and move toward favorable chemical gradients confers a strong fitness advantage. Flagella-driven motility allows bacteria to navigate toward nutrient hotspots and away from unfavorable conditions, facilitating rapid colonization of resource-rich microenvironments [64]. In *L. welshimeri*, we observed that flagellar genes were targeted by positive selection, which likely contributed to its niche expansion through enhanced motility and chemotaxis. Consistent with this finding, laboratory and evolutionary experiments have shown that, under selective pressure, *Escherichia coli* can evolve enhanced swimming velocity and altered tumbling rates, improving habitat colonization and nutrient acquisition [65]. In addition, the increased diversity of pathways in cellular processes and environmental information processing observed in *L. welshimeri* may reflect an expanded repertoire of sensing and regulatory systems (e.g., phosphotransferase system and two-component systems), and together with the presence of flagellar assembly pathways, enhance its ability to detect and respond to diverse external stimuli, benefiting niche expansion.

*L. welshimeri* also appears to leverage flagella to expand its ecological niche and broaden its geographic range by enhancing host attachment and colonization, thereby enabling frequent dispersal across landscapes. Flagellar genes can facilitate adhesion, host attachment, and colonization [62, 66–68]. For example, in *Clostridioides difficile*, flagellin is essential for initial attachment to mucosal surfaces [67], while in *Helicobacter pylori*, deletion of *fliK* reduces adhesion to gastric mucosal cells [68]. Both *H. pylori* and *Campylobacter jejuni* regulate flagellar biosynthesis and chemotaxis to penetrate viscous milieus, such as gastrointestinal mucus, facilitating host colonization [62]. In *Listeria*, flagella have also been shown to enhance host cell adhesion and invasion by increasing the bacterium-host contact frequency and facilitating early attachment [69–71]. Such host interactions can help bacteria overcome geographic barriers and promote dispersal over large spatial scales [72]. Indeed, in this study, we observed minimal dispersal limitation in *L. welshimeri*, as indicated by a homogenized population structure across geographic locations. Similar patterns have been reported in *L. monocytogenes* lineage I, *Vibrio parahaemolyticus,* and *V. fluvialis,* where frequent movement facilitates gene flow and limits population divergence [53, 73, 74]. Further, using spatial phylogenetics, we identified wild birds as a plausible dispersal vehicle for *L. welshimeri*. Wild birds, such as starlings, pigeons, and migratory gulls, have been reported as carriers and dispersal vectors of *Listeria* [31, 75, 76]. Avian-mediated dispersal has also been shown to shape the biogeographic patterns of *L. monocytogenes* [75] and *E. coli* [77].

By nature, non-motile bacteria face intrinsic constraints on niche expansion because they lack motility. However, based on our systematic, large-scale survey, the non-motile species *L. booriae* was found to be highly ecologically successful in soil, occupying a broad range of environmental conditions [28]. In this study, we show that the broad geographic range of *L. booriae* is underpinned by extensive metabolic versatility enabled by a highly flexible pangenome for environmental adaption. Specifically, its pangenome is characterized by a diverse repertoire of genes involved in metabolic pathways. The metabolic network of *L. booriae* may have evolved greater complexity and connectivity as its pangenome expanded. This expansion likely facilitates the exploitation of diverse ecological resources during niche colonization [9]. Also, genes involved in metabolic functions, spanning both the core and accessory genome, show strong signatures of positive selection in *L. booriae*. Major adaptive traits included inorganic ion transport, amino acid and coenzyme metabolism, and protein biosynthesis. These processes are essential for nutrient acquisition, enzymatic activity, and cellular maintenance [78]. Enhanced metabolic efficiency, flexibility, and diversification likely enable *L. booriae* to thrive across nutrient-variable environments, compensating for its lack of motility and limited dispersal capacity.

Our data suggest that key environmental pressures that may shape the pangenome variation and metabolic versatility of *L. booriae* include climatic factors and microbial interactions. Consistent with this result, genes involved in metabolism in soil microbiomes showed strong associations with climatic variation [79–81]. Precipitation, in particular, functions as a dominant selective filter by alternately promoting the solubilization of essential mineral ions and nutrients, such as Fe and amino acids, and driving their depletion [82, 83]. Our results suggest that *L. booriae* responds to these moisture-driven fluctuations through the retention or positive selection of genes involved in inorganic ion, coenzyme, and amino acid transport and metabolism, including high-affinity scavenging and utilization systems for Fe, such as enterochelin esterase, Fe ABC transporters, and siroheme synthase, as well as amino acid transport and metabolism genes, including members of the major facilitator superfamily and pyridoxal phosphate dependent aminotransferases. Similar adaptive strategies have been reported in rhizosphere communities, where Fe metabolism genes are enriched in Actinobacteria under drought conditions that reduce Fe bioavailability [81]. Furthermore, because cofactor scarcity can drive compensatory amino acid uptake to maintain cellular homeostasis [84], the concurrent selection across these functional categories in *L. booriae* suggests metabolic flexibility that likely enables efficient resource scavenging under nutrient limitation while minimizing enzymatic constraints when resources are more abundant, as observed in *E. coli* [85].

In addition to abiotic influences, we identified Proteobacteria, Chloroflexi, and Actinobacteria as bacterial taxa most strongly associated with pangenome flexibility in *L. booriae*, particularly for genes involved in inorganic ion transport and metabolism. Members of these phyla are known to possess diverse systems for inorganic ion acquisition and nutrient transformation [86, 87], suggesting that community composition may exert selective pressures on genes related to these functions in *L. booriae*. For example, *Listeria* does not synthesize siderophores for Fe acquisition but can utilize a range of exogenous organic compounds, including siderophores and catechol ligands, to support growth under iron-limited conditions [88, 89], raising the possibility that *L. booriae* relies, at least in part, on co-occurring bacteria to provide extracellular compounds that facilitate the acquisition of essential inorganic ions.

Together, our findings delineate distinct genomic and ecological strategies underlying the broad geographic ranges of motile and non-motile *Listeria* species, refining our understanding of how bacteria with contrasting lifestyles achieve broad ecological distributions and resilience to environmental pressures. Notably, our study highlights often-overlooked mechanisms that enable non-motile bacteria to persist and diversify across ecosystems. These insights have important implications for controlling motile and non-motile pathogens and for minimizing environmental transmission. Given the importance of *Listeria* to food safety, and the close monitoring of multiple species beyond the pathogen *L. monocytogenes* in the food industry, our findings also inform risk assessment and targeted interventions throughout food production, processing, and storage.

## Data availability

Whole-genome sequencing data for 141 *L. welshimeri* and 90 *L. booriae* soil isolates used in this study have been submitted to the National Center for Biotechnology Information’s (NCBI) BioProject database under accession number PRJNA561882, along with associated environmental metadata, previously published in Liao et al. (2021) [28]. The 16S rRNA gene amplicon sequencing data analyzed in this study are available under BioProject accession number PRJNA749132, as published in Liao et al. (2023) [39]. Trimmed paired-end reads for the nine *L. welshimeri* wild bird isolates used in this study have been deposited in the NCBI Sequence Read Archive (SRA), and their assembled genomes are available in NCBI GenBank under the accession numbers listed in **Supplementary Table 1**. All data needed to evaluate the conclusions in the paper are present in the paper and/or the Supplementary Information.

## Code availability

Code to replicate all analyses is publicly available at https://github.com/leaph-lab/Listeria_motility_eco_MS.

## Acknowledgements

We are grateful to Dr. Renato Orsi for assistance in identifying *L. welshimeri* wild bird isolates, and to the members of LEAPH (the Liao Laboratory) for their enriching discussions. Part of this work was funded by the Virginia Tech Graduate and Professional Student Senate Graduate Research Development Program (Y.-X.G.) and 4-VA Collaborative Research Grant (J.L.).

## Author contributions

Conceptualization, J.L.; Data Curation, Y.-X.G., K.J.C., M.W., J.L.; Investigation, Y.-X.G., S.H., J.L.; Formal Analysis, Y.-X.G., J.L.; Visualization, Y.-X.G.; Writing – original draft preparation, Y.-X.G.; Writing – review & editing, K.J.C., M.W., J.L.; Supervision, J.L.

