## Supplementary material for "Motile and non-motile *Listeria* species adopt distinct genomic and ecological strategies to achieve broad geographic ranges in soil": Supplmentary Information

### Supplementary Figures

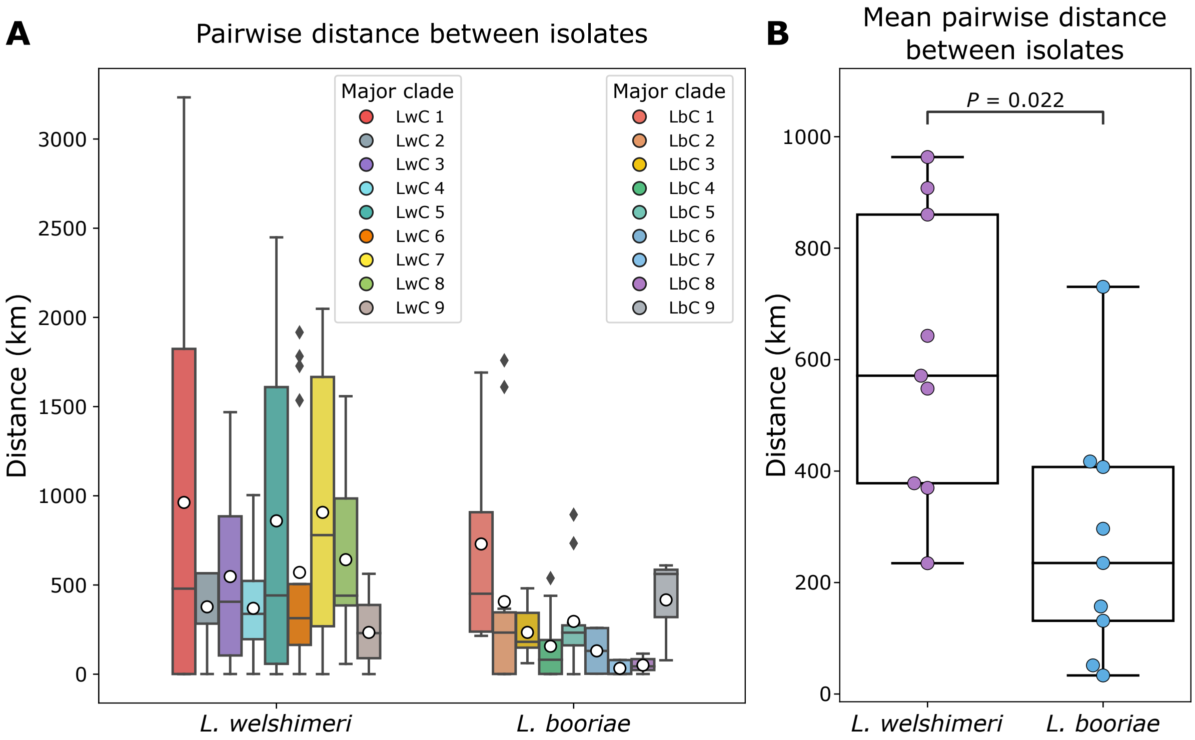


**Supplementary Figure 1. Geographic distribution of the major clades of *L. welshimeri* and *L. booriae*. (A)** Pairwise geographic distances between isolates within major clade of each species. Major clades were identified based on the phylogenetic trees shown in **Fig.1 C-D** and are color-coded, with "Lw" and "Lb" indicating *L. welshimeri* and *L. booriae*, respectively. White dots indicate the mean pairwise distance within each major clade. **(B)** Mean pairwise geographic distance between isolates across major clades compared between *L. welshimeri* and *L. booriae*. Two-sided Mann-Whitney (MW) *U P* value is annotated. For (**A**) and (**B**), box plots show the interquartile range (IQR), with the median indicated by a horizontal line and whiskers extending to 1.5 times IQR.

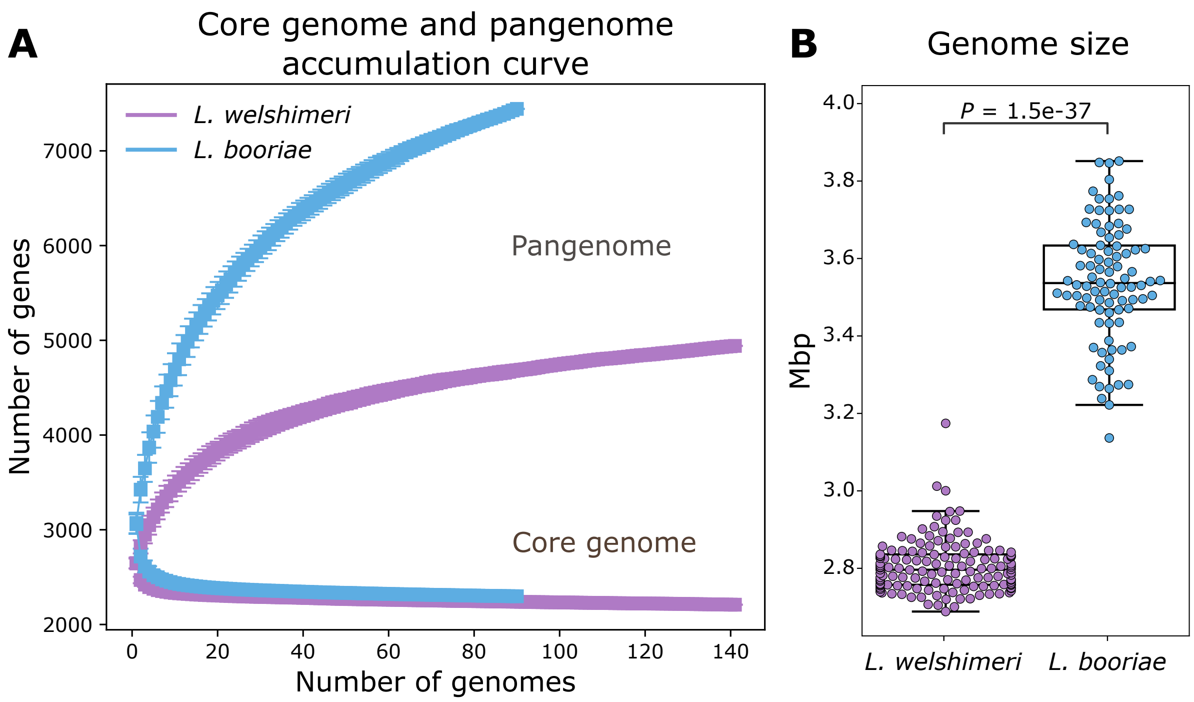


**Supplementary Figure 2. Pangenome openness and genome size of *L. welshimeri* and *L. booriae*. (A)** Accumulation curves of the core genome and pangenome of *L. welshimeri* and *L. booriae*. The lower curves represent core genome accumulation, whereas the upper curves represent pangenome accumulation. These curves were adapted from **Extended Data Fig. 7** in Liao et al (2021) [1]. **(B)** Genome sizes of isolates compared between *L. welshimeri* and *L. booriae*. Box plots show the IQR, with the median indicated by a horizontal line and whiskers extending to 1.5 times IQR. Two-sided MW *U P* value is annotated.

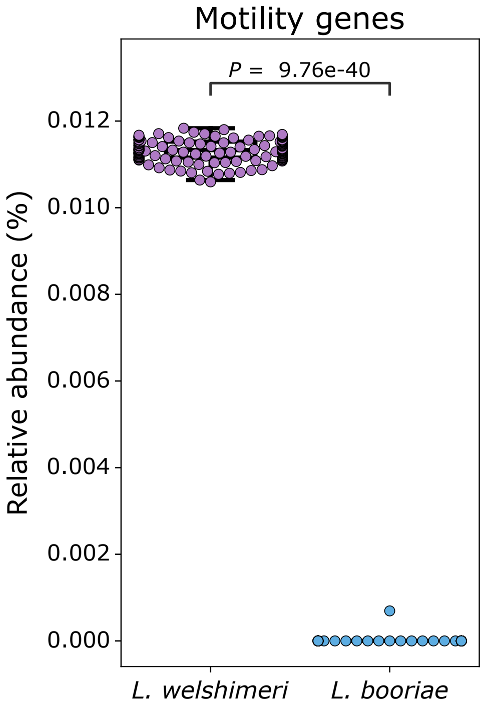


**Supplementary Figure 3. Relative abundance of putatively functional motility genes compared between *L. welshimeri* and *L. booriae*.** Box plots show the IQR, with the median indicated by a horizontal line and whiskers extending to 1.5 times IQR. Two-sided MW *U P* value is annotated.

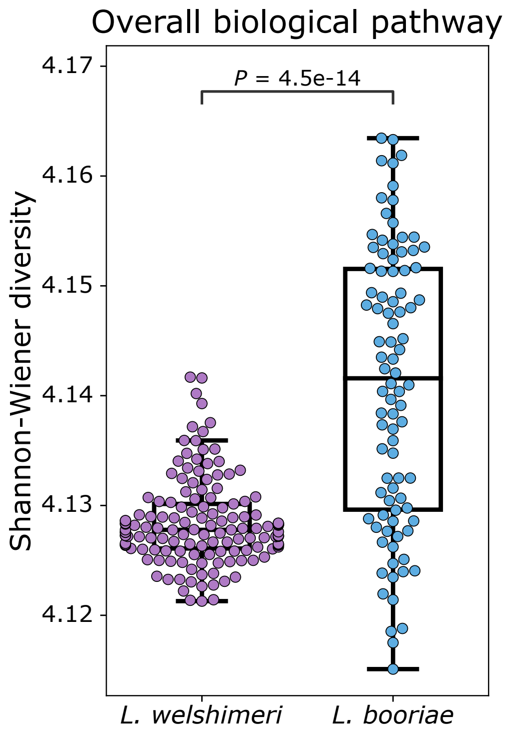


**Supplementary Figure 4. Shannon-Wiener diversity of overall biological pathways compared between *L. welshimeri* and *L. booriae*.** Box plots show the IQR, with the median indicated by a horizontal line and whiskers extending to 1.5 times IQR. Two-sided MW *U P* value is annotated.

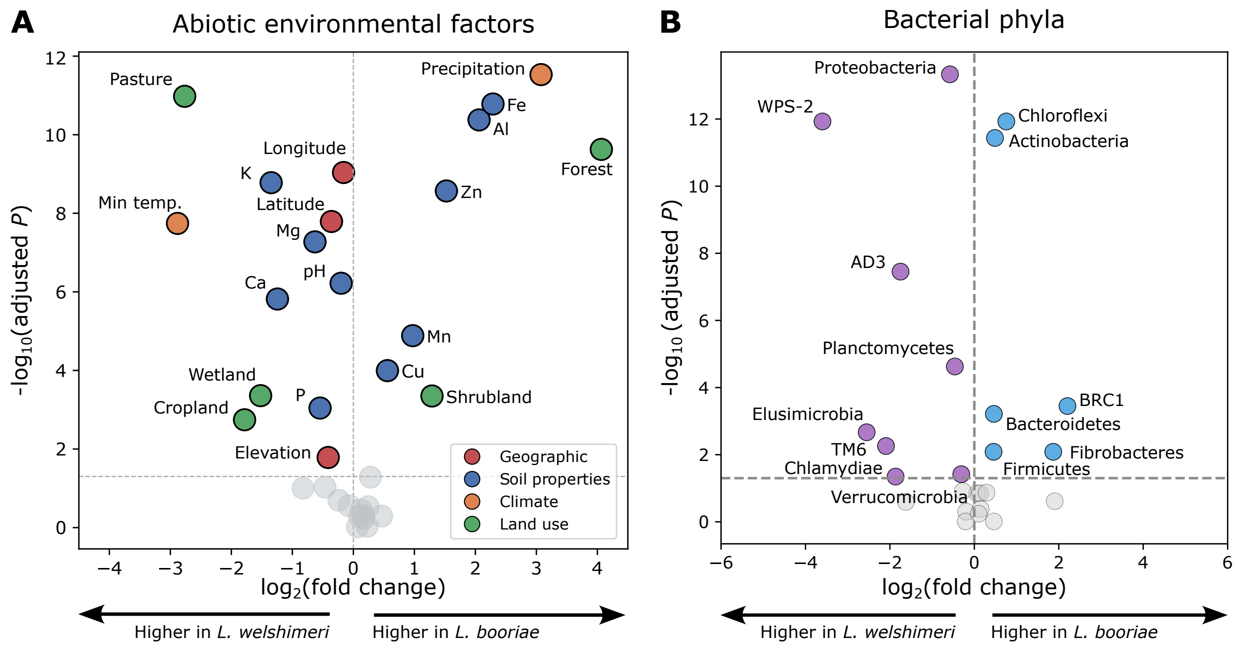


**Supplementary Figure 5. Individual abiotic and biotic environmental factors compared between *L. welshimeri* and *L. booriae*. (A-B)** Volcano plots showing fold change (x-axis) versus statistical significance (y-axis, adjusted two-sided MW *U* test) for **(A)** abiotic factors and **(B)** relative abundance of bacterial phyla. Points above the gray dashed line indicate adjusted $P < 0.05$ and are color-coded by abiotic factor groups in (**A**) and by species with a higher relative abundance in (**B**). Abbreviations of abiotic factors are described in Methods. WPS-2, AD3, TM6, and BRC1 represent candidate bacterial phyla that remain uncultured under laboratory conditions.

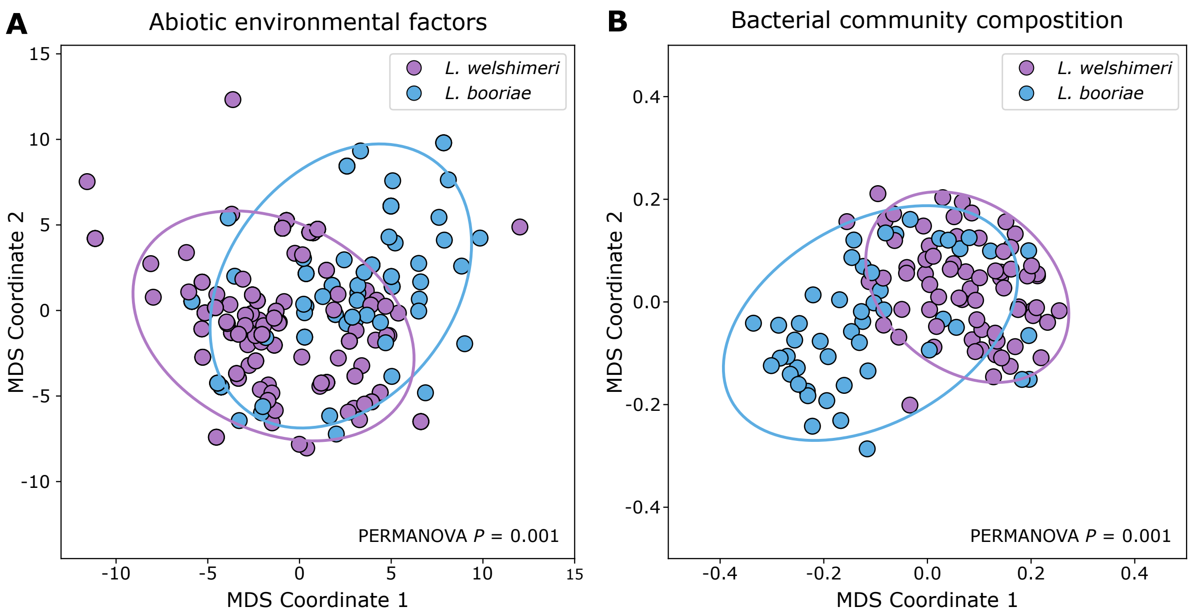


**Supplementary Figure 6. Overall abiotic and biotic environmental conditions compared between *L. welshimeri* and *L. booriae*.** **(A-B)** Multidimensional scaling (MDS) analysis based on **(A)** Euclidean distances of abiotic factors and **(B)** weighted UniFrac distances of bacterial community composition derived from OTUs. Points represent isolates and are color-coded by species. Ellipses indicate two standard deviations from the mean. PERMANOVA $P < 0.05$ denotes significant clustering by species.

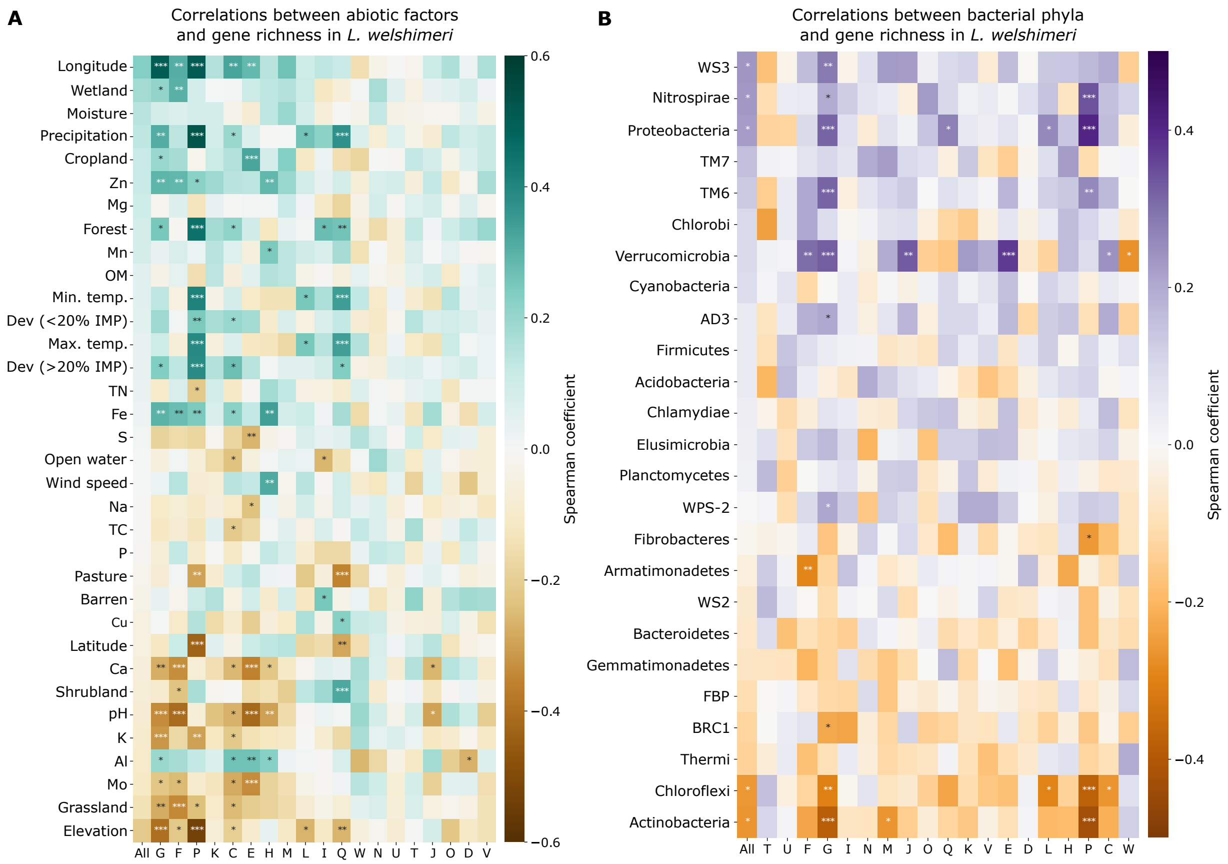


**Supplementary Figure 7. Associations between gene richness in *L. welshimeri* and abiotic environmental factors and bacterial community composition. (A-B)** Spearman's correlations between **(A)** abiotic factors and gene richness, both overall and for COGs, and **(B)** the relative abundance of bacterial phyla and gene richness, both overall and for COGs, in *L. welshimeri*. Abiotic factors and bacterial phyla are ordered by the descending correlation coefficients of overall ("All") gene richness. Positive and negative correlations are shown in green and brown for abiotic factors, and in purple and orange for bacterial phyla, respectively. Abbreviations of COGs and abiotic factors are described in figure legend of **Fig. 2** and in the Methods, respectively. TM6, WS3, WS2, AD3, WPS-2, FBP, BRC1, and TM7 represent candidate bacterial phyla that remain uncultured under laboratory conditions.

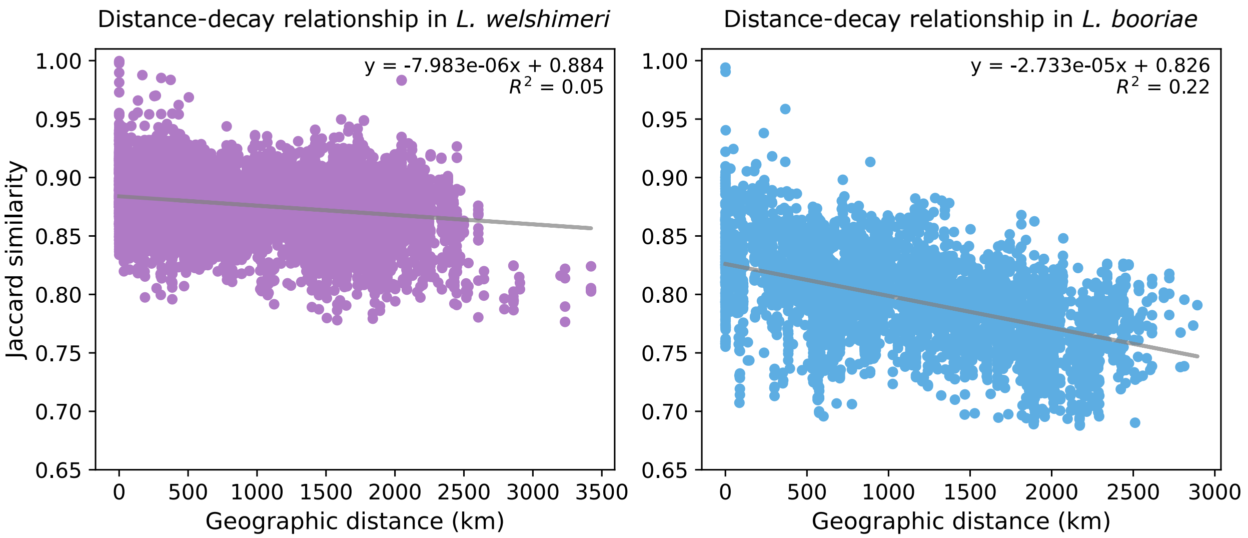


**Supplementary Figure 8. Dispersal patterns of *L. welshimeri* and *L. booriae*.** Distance-decay relationship in *L. welshimeri* and *L. booriae* was inferred by the linear regression for genetic similarities measured by Jaccard similarity of gene absence/presence and geographic distances. A steeper negative slope with a higher $R^2$ indicates a stronger distance-decay relationship.

**Supplementary Tables for this manuscript include the following:**

**Supplementary Table 1.** Quality statistics of nine *L. welshimeri* wild bird isolates for whole-genome sequencing.

**Supplementary Table 2.** List of reference motility and chemotaxis genes retrieved from BIGSdb-*Lm* for BLASTN searches.

**Supplementary Table 3**. List of KEGG pathway identifiers used in pathway diversity analysis.

**Supplementary Table 4.** List of independent variables with variance inflation factors (VIF) below 10 used in the variation partitioning analysis (VPA), along with their corresponding model and variable categories.

**Supplementary Table 5.** List of genes under positive selection without evidence of homologous recombination.
